# Gi-Coupled Receptor Activation Potentiates Piezo2 Currents via Gβγ

**DOI:** 10.1101/722637

**Authors:** John Smith Del Rosario, Yevgen Yudin, Songxue Su, Cassandra Hartle, Tooraj Mirshahi, Tibor Rohacs

## Abstract

**SUMMARY:** Dysregulation of mechanosensitive Piezo2 channels is a hallmark of mechanical allodynia, yet the cellular mechanisms that sensitize mechanoreceptors are still poorly understood. Activation of Gi-coupled receptors sensitizes Piezo2 currents, but whether Gi-coupled receptors regulate the activity of Piezo2 channels is not known. Here, we found that activation of Gi-coupled receptors potentiates Piezo2 currents in dorsal root ganglion (DRG) neurons and in heterologous systems, but inhibits Piezo1 currents in heterologous systems. The potentiation, or inhibition of Piezo currents is abolished when blocking Gβγ with the c-terminal domain of the beta-adrenergic kinase (βARKct). Pharmacological inhibition of kinases downstream of Gβγ, phosphoinositide 3-kinase (PI3K) and mitogen-activated protein kinase (MAPK), also abolished the potentiation of Piezo2 currents. Hence, our studies illustrate an indirect mechanism of action of Gβγ to sensitize Piezo2 currents after activation of Gi-coupled receptors.

## INTRODUCTION

Our ability to sense environmental cues is vital for our daily activities. Our tactile world relies on a wide variety of mechano-sensory neurons that innervate our skin and are responsible for sensing innocuous and painful stimuli attributed to low-threshold mechanoreceptors (LTMRs) and nociceptors, respectively. These neurons detect external mechanical stimulation and convert it into ionic currents using mechano-gated ion channels, including the excitatory Piezo2 non-selective cation channels and the inhibitory K^+^ selective TREK1 and TREK2 channels (Alloui et al., 2006; Coste et al., 2010). Piezo2 channels are intrinsically gated by force and mediate rapidly-adapting mechanically activated (RA-MA) currents in DRG neurons and when heterologously expressed. Reports have shown that Piezo2 channels play a pivotal role in the detection of light touch, proprioception and mechanical pain in mice and humans (Chesler et al., 2016; Eijkelkamp et al., 2013; Mahmud et al., 2017; Murthy et al., 2018; Ranade et al., 2014; Szczot et al., 2018; Woo et al., 2015). In addition, these channels are essential regulators of embryonic development, lung expansion, reverse polarity currents of auditory hair cells, Merkel cell mechanotransduction, baroreception and itch (Alisch et al., 2017; Coste et al., 2013; Feng et al., 2018; Nonomura et al., 2017; Wu et al., 2017; Zeng et al., 2018). Despite the substantial progress in describing the physiological importance of Piezo2, little is known about how endogenous signaling pathways modulate the activity of these channels.

DRG neurons express a wide variety of G-Protein coupled receptors. Gs- and Gq-coupled receptors generally increase excitability of DRG neurons, and many of them are activated by inflammatory mediators such as prostaglandins and bradykinin. Activation of the Gq-coupled bradykinin receptor B2 was shown to sensitize Piezo2 currents in heterologous systems and in DRG neurons through a mechanism involving the activation of PKA and PKC, an effect that may play a role in inflammatory mechanical allodynia (Dubin et al., 2012).

Whether the activation of Gi-coupled receptors influence the activity of Piezo2 channels has not been studied yet. Activation of Gi-coupled receptors leads to inhibition of adenylate cyclase, activation of G-protein coupled inwardly rectifying K^+^ (GIRK) channels and inhibition of voltage-gated Ca^2+^ channels (VGCC) and of transient receptor potential melastatin 3 (TRPM3) channels in DRG neurons through direct binding of Gβγ (Badheka et al., 2017; Dembla et al., 2017; Quallo et al., 2017; Reuveny et al., 1994; Robertson and Taylor, 1986). In addition, Gβγ is also known to induce the activation of phosphoinositide 3-kinase gamma (PI3Kγ) and mitogen-activated protein kinase (MAPK) (Clapham and Neer, 1997; Khan et al., 2013; Lopez-Ilasaca et al., 1997; Touhara et al., 1995).

Many different Gi-coupled receptors, such as GABA_B_ and opioid receptors are expressed in DRG neurons; their activation generally reduces excitability, and exert analgesic effects (Yudin and Rohacs, 2018). Activation of many Gi-coupled receptors however was shown to induce mechanical hypersensitivity, the mechanism of which is not understood (Araldi et al., 2016, 2018; Yudin and Rohacs, 2018)

Here we found that activation of Gi-coupled receptors potentiated native Piezo2-mediated currents in DRG neurons and heterologous expressed Piezo2 channels in HEK293 cells without affecting the number of channels at the plasma membrane. Surprisingly, the currents of the closely related Piezo1 channels were inhibited after activation of Gi-coupled receptors in HEK293 cells. The potentiation Piezo2 and inhibition of Piezo1 channels were abolished when blocking Gβγ. Furthermore, inhibition of PI3K and MAPK also abolished the potentiation of Piezo2 currents. In conclusion, our data show that Gi-coupled receptors potentiate Piezo2 currents through a mechanism potentially involving a Gβγ-dependent PI3K and MAPK activation.

## RESULTS

### GABA_B_ Receptor Activation Potentiates Piezo2-like Currents in DRG Neurons

At the RNA level, GABA_B_ receptors are the highest Gi-coupled receptors expressed in DRG neurons (Thakur et al., 2014; Usoskin et al., 2015). Therefore, we tested if activation of GABA_B_ receptors could influence the activity of Piezo2 channels. We mechanically stimulated, with a glass probe, large diameter DRG neurons (cell capacitance >40 pF), which are most likely mechanoreceptors (Narayanan et al., 2018), and recorded rapidly adapting-mechanically activated (RA-MA) currents, attributed to Piezo2 channels (Coste et al., 2010; Woo et al., 2015). First, we repetitively applied short mechanical stimulations of the same amplitude every 30 seconds, and after 3 minutes of recording mechanically activated currents, we activated GABA_B_ receptors by treating the cells with 25 µM baclofen. Activation of GABA_B_ receptors potentiated RA-MA currents in 50% of neurons tested (Figure 1A-B).

**Figure 1.**
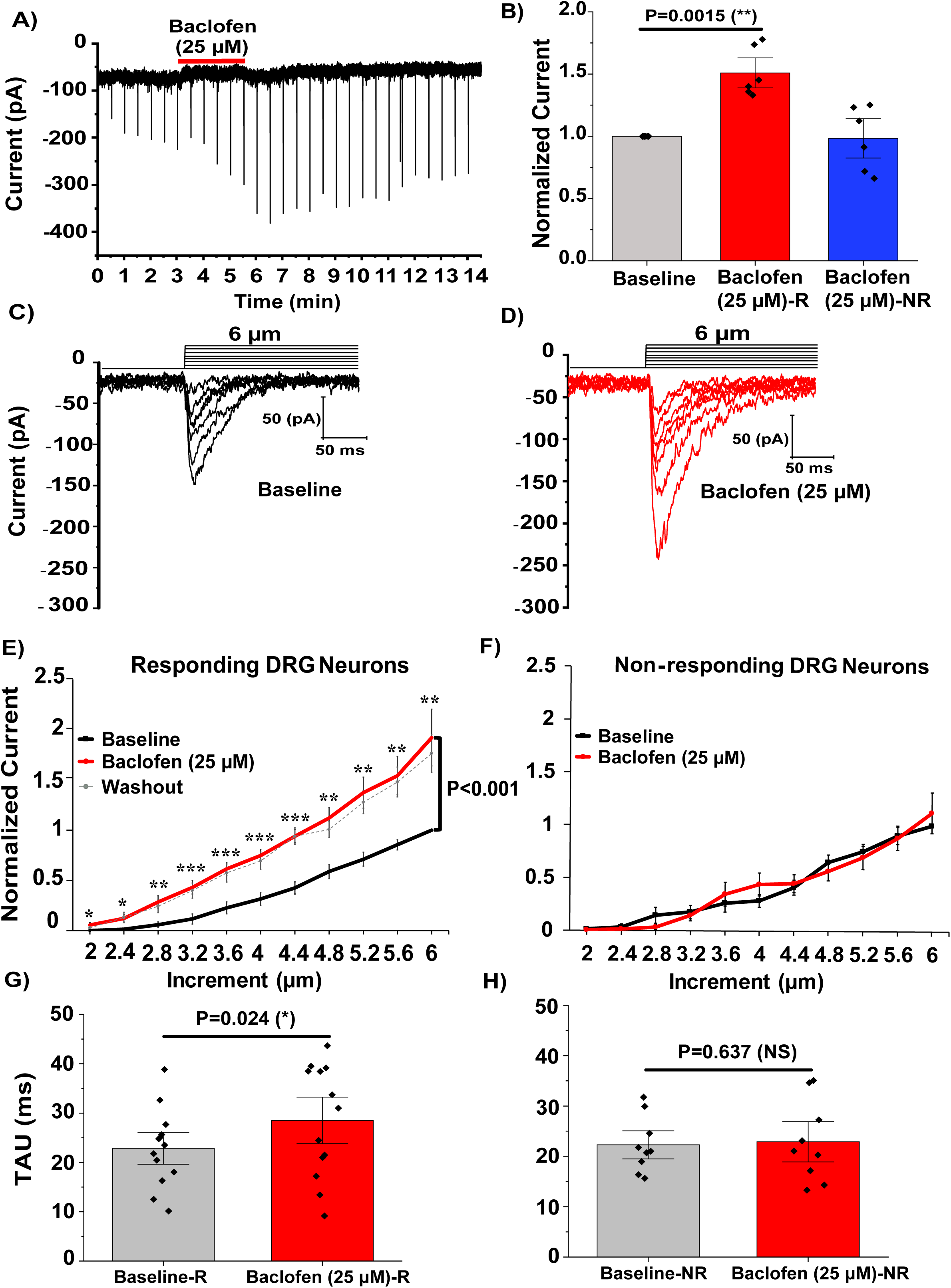
GABA_B_ Receptor Activation Potentiates Piezo2-like Currents in DRG Neurons. (A) Representative of RA-MA currents recorded in whole cell patch clamp experiments at −60 mV in large DRG neurons in response to mechanical indentation with a blunt glass probe displaced at 6.8 µm (continuous protocol) before (black downwards spikes) and after (red line) exposure to 25 µM baclofen. The downwards spikes represent individual MA currents induced by repetitive mechanical stimuli every 30 s for 200 ms stimulus duration. (B) Quantification of normalized RA-MA currents of responding DRG neurons (R, red) (n=6, **P<0.01, Paired t-test) and non-responding DRG neurons (NR, blue, n=6). Data are shown as mean ± SEM. (C-D) Representative of RA-MA currents recorded at −60 mV in large DRG neurons in response to repetitive mechanical stimulation with a blunt glass probe displaced 2-6 µm every 15 s (step protocol) before (black) and after (red) exposure to 25 µM Baclofen. The mechanical step protocol that indicates the displacement of mechanical probe is also shown. (E) Quantification of normalized MA currents of responding large DRG neurons [n=13, *P<0.05, **P<0.01, ***P<0.001, before (black) and after (red) baclofen treatment. Dashed grey line shows average MA currents 3-15 minutes after washout of baclofen (not included in the statistical analysis). Repeated measures ANOVA with student’s t-test no corrections]. Data are shown as mean ± SEM. (F) Quantification of normalized MA currents of non-responding DRG neurons (n=9). (G-H) Quantification of MA current inactivation rate (Tau) measured at 6 µm of responding (R) large DRG neurons (n=13, *P<0.05, Paired t-test) and non-responding large (NR) DRG neurons (n=9,NS,not significant) before (gray) and after (red) baclofen application. Data are shown as mean ± SEM.

We also measured RA-MA currents using a 2-6 µm step protocol, in which we increased the displacement of the glass probe every 15 s thus increasing the probability of channel opening. After recording the basal RA-MA currents, the cells were treated with 25 µM baclofen for 3 mins and RA-MA currents were re-recorded for comparison. As expected, activation of GABA_B_ receptors also potentiated Piezo2 currents almost 2-fold (Figure 1C-E), in 59 % of the neurons, and had no effect in 41%% of DRG neurons tested (Figure 1F) probably because not every DRG neuron co-expresses GABA_B1_ and GABA_B2_ receptors and the obligatory heterodimerization of these receptors is required for proper G-protein signaling (Padgett and Slesinger, 2010; Yudin and Rohacs, 2018). In both protocols, baclofen-induced potentiation was long lasting and persisted for several minutes after removal of the agonist (Fig. 1A and Fig. 1E, dashed line). We also noticed that the kinetics of inactivation of responding cells (R) were somewhat slower after treating the cells with baclofen (Figure 1G).

### GABA_B_ Receptor Activation Potentiates Piezo2 Currents in Heterologous Systems

DRG neurons are a heterogenous population of cells that express a variety of MA ion channels with distinct kinetics of inactivation: rapidly adapting (RA), intermediate adapting (IA) and slowly adapting (SA) (Borbiro et al., 2015; Prato et al., 2017). To determine if the potentiation of Piezo2 currents after baclofen treatment was specific to Piezo2 channels, we co-expressed Piezo2 and GABA_B_ receptors in HEK293 cells with GFP (as a marker for visualization). Using the same step protocol as in DRG neurons, we recorded Piezo2 currents and compared them before and after application of baclofen. Activation of GABA_B_ receptors by baclofen (25 µM) also potentiated Piezo2 currents in HEK293 cells (Figure 2A-C). Furthermore, there was no statistically significant difference in the kinetics of inactivation of Piezo2 currents measured after baclofen application (Figure 2D). In addition, Piezo2 currents recorded from HEK293 cells transfected with only Piezo2 channels were not potentiated after baclofen treatment, showing that the activation of GABA_B_ receptor was required (Figure 2E).

**Figure 2.**
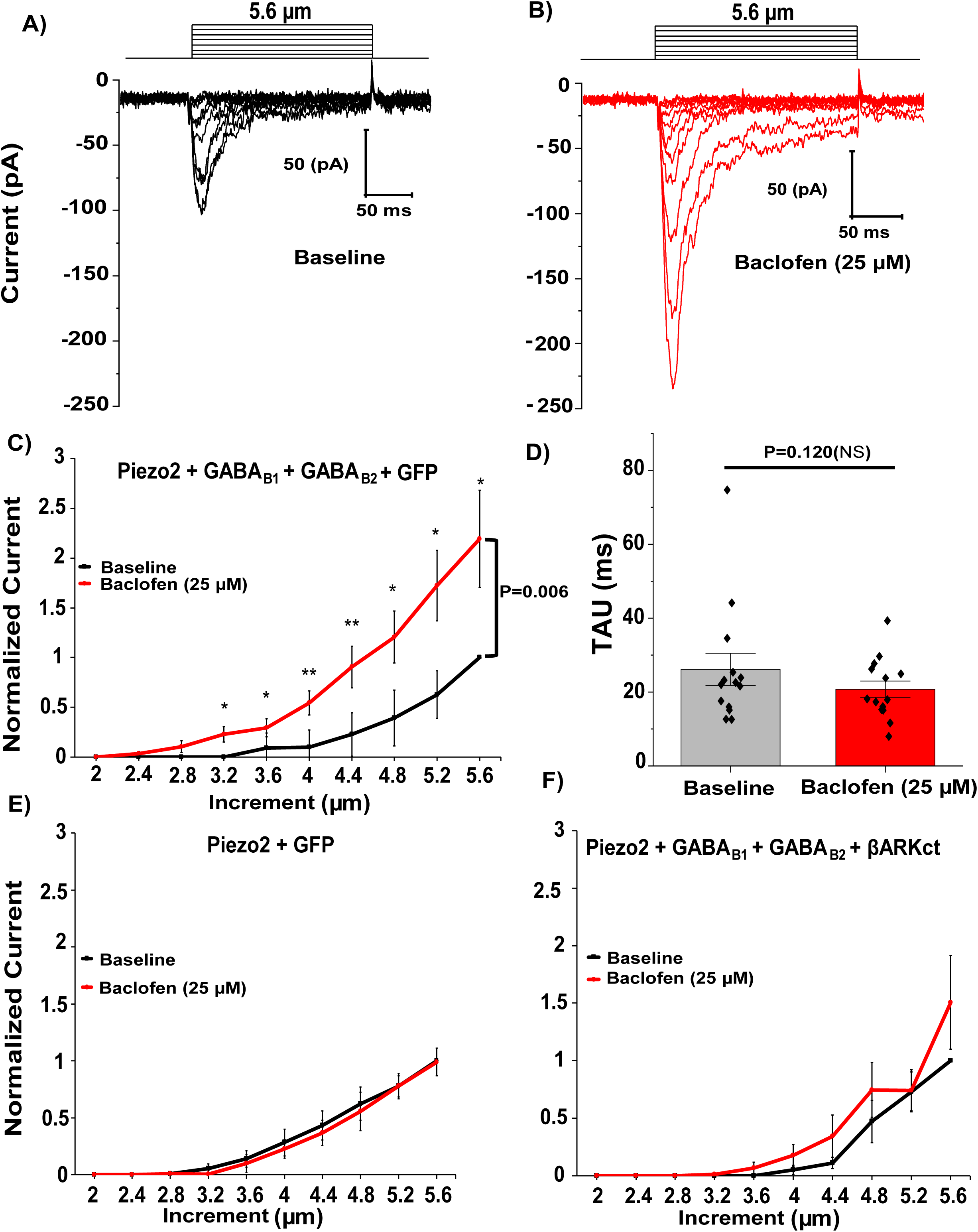
GABA_B_ Receptor Activation Potentiates Piezo2 Currents in a Heterologous System via Gβγ. (A-B) Representative MA currents recorded in whole cell patch clamp experiments at −60 mV in HEK293 cells transiently transfected with Piezo2 ^+^GABA_B1_^+^ GABA_B2_ ^+^ GFP (for visualization) in response to repetitive mechanical stimulation with a blunt glass probe displaced 2-5.6 µm every 15 s (step protocol) before (black) and after (red) exposure to 25 µM baclofen. The mechanical step protocol that indicates the displacement of mechanical probe is also shown. (C) Quantification of normalized MA currents of HEK293 cells transfected with Piezo2^+^GABA_B1_^+^ GABA_B2_ ^+^ GFP [C, n=14, *P<0.05, **P<0.01, before (black) and after (red) baclofen treatment, Repeated measures ANOVA with student’s t-test no corrections]. (D) Quantification of MA current inactivation rate (Tau) measured at indentation of 5.6 µm of HEK293 cells transfected with Piezo2^+^GABA_B1_^+^ GABA_B2_ ^+^ GFP before (gray) and after (red) baclofen application (n=14, NS, not significant; Paired t-test) (E-F) Quantification of normalized MA currents of HEK293 cells transfected Piezo2 ^+^ GFP (E, n=7) and Piezo2^+^GABA_B1_^+^ GABA_B2_ ^+^ βARKct-mCherry (F, n=8) before (black) and after (red) application of 25 µM baclofen. Data are shown as mean ± SEM.

### Potentiation of Piezo2 Currents after activation of GABA_B_ Receptors is via Gβγ

Activation of Gi-coupled receptors leads to dissociation of the G-proteins: Gαi and Gβγ. The Gαi subunit inhibits adenylate cyclase, while Gβγ inhibits neuronal activity through activation of GIRK channels and VGCC (Logothetis et al., 1987). To test whether Piezo2 inhibition requires Gβγ, we co-transfected HEK293 cells with Piezo2 channels, GABA_B_ receptors and the c-terminal domain of the beta adrenergic kinase (βARKct), which is known to block the effects of Gβγ (Badheka et al., 2017; Logothetis et al., 1987; Sato et al., 2015). Cotransfection of βARKct abolished the potentiation of Piezo2 currents after activation of GABA_B_ receptors by baclofen (Figure 2F), indicating the involvement of Gβγ.

### Activation of other Gi-coupled Receptors also Potentiates Piezo2 currents in DRG neurons and Heterologous Systems

Our data show that activation of GABA_B_ receptors potentiate Piezo2 currents in DRG neurons and in HEK293 cells. Next we tested whether this potentiation was caused by a general Gi-coupled receptor mechanism, or one specific to GABA_B_ receptors, as it was reported for TRPV1 channels (Hanack et al., 2015). We transfected HEK293 cells with Piezo2 channels and the Gi-coupled D2 dopamine receptors, and we found that the D2/D3 agonist quinpirole (200 nM) potentiated Piezo2 currents (Figure 3A). On the other hands, cells transfected with only Piezo2 channels did not show any MA current potentiation after quinpirole treatment (Figure 3B), indicating the role of D2 receptor activation.

**Figure 3.**
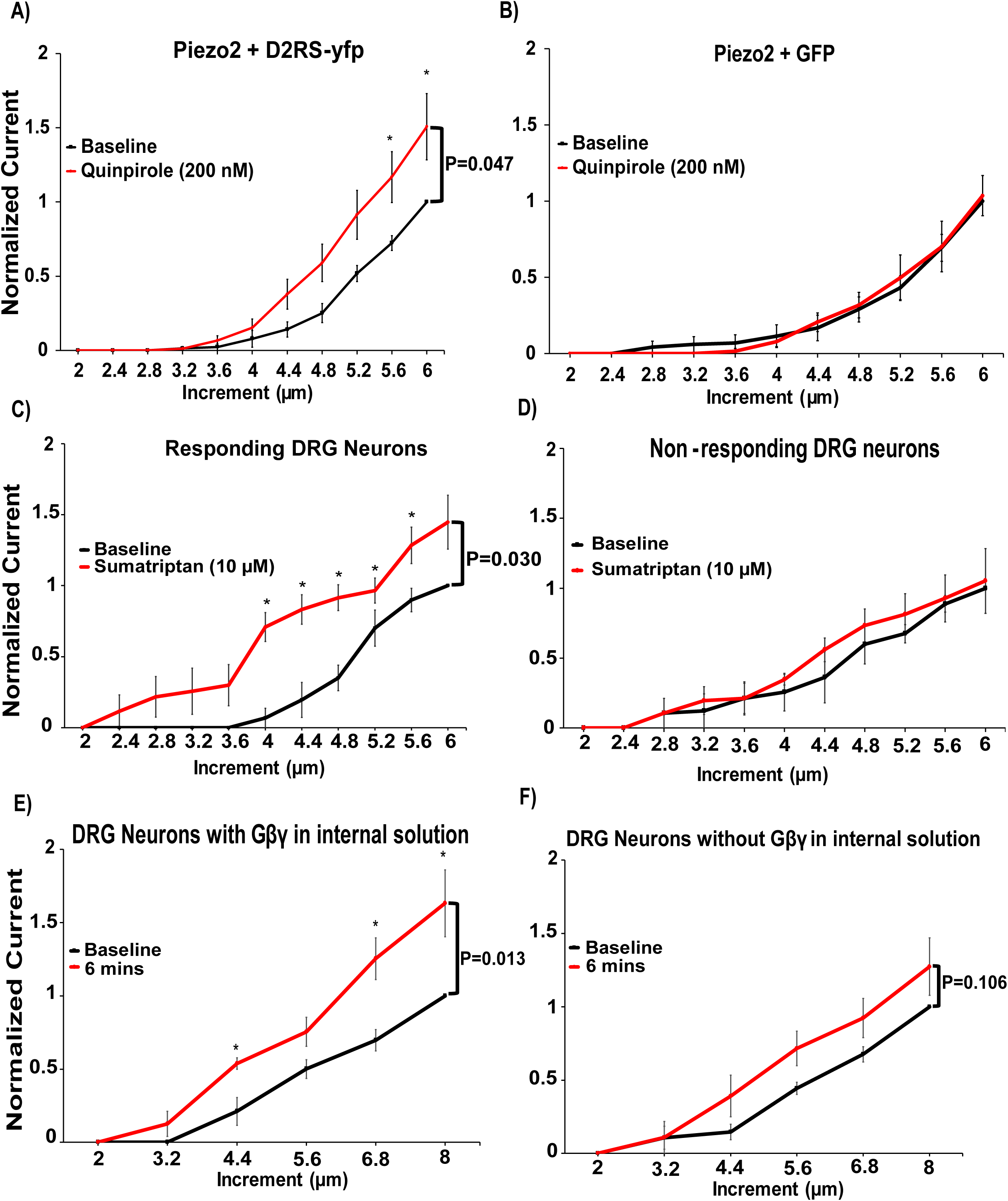
Activation of other Gi-coupled Receptors also Potentiates native and recombinant Piezo2 currents. (A-B) Quantification of normalized RA-MA Piezo2 currents in HEK293 cells transiently transfected with Piezo2^+^D2RS-yfp (A, n=14) and HEK293 cells transfected with Piezo2^+^GFP (B, n=5) before (black) and after (red) application of quinpirole 200 nM. *P<0.05, Repeated measures ANOVA with student’s t-test no corrections. Data are shown as mean ± SEM. (C-D) Quantification of normalized RA-MA currents in responding large DRG neurons (C, n=5) versus non-responding DRG neurons (D, n=3) before (black) and after (red) application of sumatriptan 10 µM to activate the Gi-coupled serotonin receptors 1D. *P<0.05, Repeated measures ANOVA with student’s t-test no corrections. Data are shown as mean ± SEM. (E-F) Quantification of normalized RA-MA currents recorded from large DRG neurons dialyzed with 50 ng/ml Gβγ (E) or with 0.0001% Lubrol (F). *P<0.05, Repeated measures ANOVA with student’s t-test no corrections. Data are shown as mean ± SEM.

The Gi-coupled serotonin receptor 1D (Htr1d) is also highly expressed in large diameter DRG neurons (Usoskin et al., 2015). We measured RA-MA Piezo2 currents from these DRG neurons and treated them with 10 µM of sumatriptan an agonist of Htr1d and Htr1b, a clinically used medication against migraine. Sumatriptan potentiated Piezo2 currents in 63% of DRG neurons, while in 37% of the neurons it did not (Figure 3C and D). These findings imply that the effect of Piezo2 current potentiation is a general Gi-coupled receptor-dependent mechanism.

To teste if Gβγ was sufficient to potentiate Piezo 2 channels we introduced Gβγ into the intracellular solution in the whole cell patch pipette and measured RA-MA currents in DRG neurons immediately after establishment of the whole cell mode and 6 mins after. As shown in figure 3E, RAMA currents were significantly enhanced after 6 mins in neurons dialyzed with Gβγ, but not in neurons where Gβγ was omitted from the patch pipette solution (Figure 3F). These data indicate that Gβγ alone was sufficient for the potentiation of Piezo2 currents.

### GABA_B_ Receptor Activation Inhibits Piezo1 Currents in Heterologous Systems via Gβγ

Our data show that Piezo2 currents are potentiated after activation of Gi-coupled receptors through a mechanism involving Gβγ. However, vertebrates have two types of Piezo channels: Piezo1 and Piezo2. Piezo channels share about ∼50% in protein sequence identity, possess similar biophysical properties and are known to be similarly regulated by endogenous molecules (Anderson et al., 2018; Borbiro et al., 2015; Coste et al., 2010; Coste et al., 2015). Therefore, we tested if the activation of Gi-coupled receptors could also affect the activity of Piezo1 channels. We transfected HEK293 cells with Piezo1 channels and GABA_B_ receptors. We recorded Piezo1 currents every 15 s increasing the displacement of the mechanical probe closer to the cell before and after application of baclofen. In contrast to Piezo2, activation of GABA_B_ receptors by 25 µM baclofen inhibited Piezo1 currents (Figure S1A-C). This inhibition depended on activation of GABA_B_ receptors because HEK293 cells transfected with only Piezo1 channels were not inhibited after baclofen treatment (Figure S1D). Cotransfecting βARKct abolished the inhibition of Piezo1 currents by activation of GABA_B_ receptors (Figure S1E), indicating that regulation of Piezo1 currents by activation GABA_B_ receptors was also mediated by Gβγ.

### PI3K and MAPK Participate in Potentiation of Piezo2 Currents after Activation of GABA_B_ Receptors

Figure 1A and 1E shows that the potentiation of Piezo2 currents lasted several minutes, even after the removal of the Gi-coupled receptor agonist. Gβγ mediated effects on ion channel are usually quickly reversible, therefore we tested if Gβγ modulated the activity of Piezo2 channels by a direct or indirect mechanism. We found that Gβγ did not co-immunoprecipate with either Piezo1 or Piezo2 channels (Figure S2), but showed clear co-immunoprecipitation with the Gβγ regulated TRPM3 channels, suggesting that Gβγ may not be forming a complex with Piezo2 channels after activation of Gi-coupled receptors.

A potential mechanism of potentiation of Piezo2 currents could be the activation of protein kinases by Gβγ. PI3Kγ and MAPK are downstream targets of Gβγ (Clapham and Neer, 1997; Khan et al., 2013). These kinases have been associated with inflammatory hypersensitivity, NGF-signaling and inflammation in nociceptive neurons (Leinders et al., 2014; Obata and Noguchi, 2004; Zhang et al., 2005) and are possibly involved in affecting mechanically activated ion channels (Dela Paz and Frangos, 2019; Prato et al., 2017). Therefore, we tested if these kinases contribute to the potentiation of Piezo2 currents in HEK293 cells co-transfected with Piezo2 channels and GABA_B_ receptors. We created three groups: 1) cells treated with vehicle, 2) cells pre-incubated with the PI3K inhibitor Wortmannin (50 nM) and 3) cells pre-incubated with the MAPK inhibitor of U0126 (10 µM). We measured Piezo2 currents every 15 s increasing the displacement of the mechanical probe closer to the cell before and after application of baclofen. Piezo2 currents measured in cells treated with either PI3K or MAPK inhibitors were not potentiated after activation of GABA_B_ receptors, but Piezo2 currents in cells treated only with vehicle were potentiated (Figure 4A-C). These findings suggest that the prolonged potentiation of Piezo2 currents upon activation of Gi-coupled receptors is caused by activation of Gβγ-downstream targets: PI3K and MAPK. PI3K activation was shown to increase surface expression of TRPV1 channels (Zhang et al., 2005), therefore we also tested if activation of GABA_B_ receptors induces a change in Piezo2 surface expression total internal reflection fluorescence (TIRF) microscopy. Figure S3 shows that the plasma membrane fluorescence of the GFP tagged Piezo2 did not change in response to baclofen activation in HEK cells transfected with Piezo2-GFP and GABA_B_ receptors.

**Figure 4.**
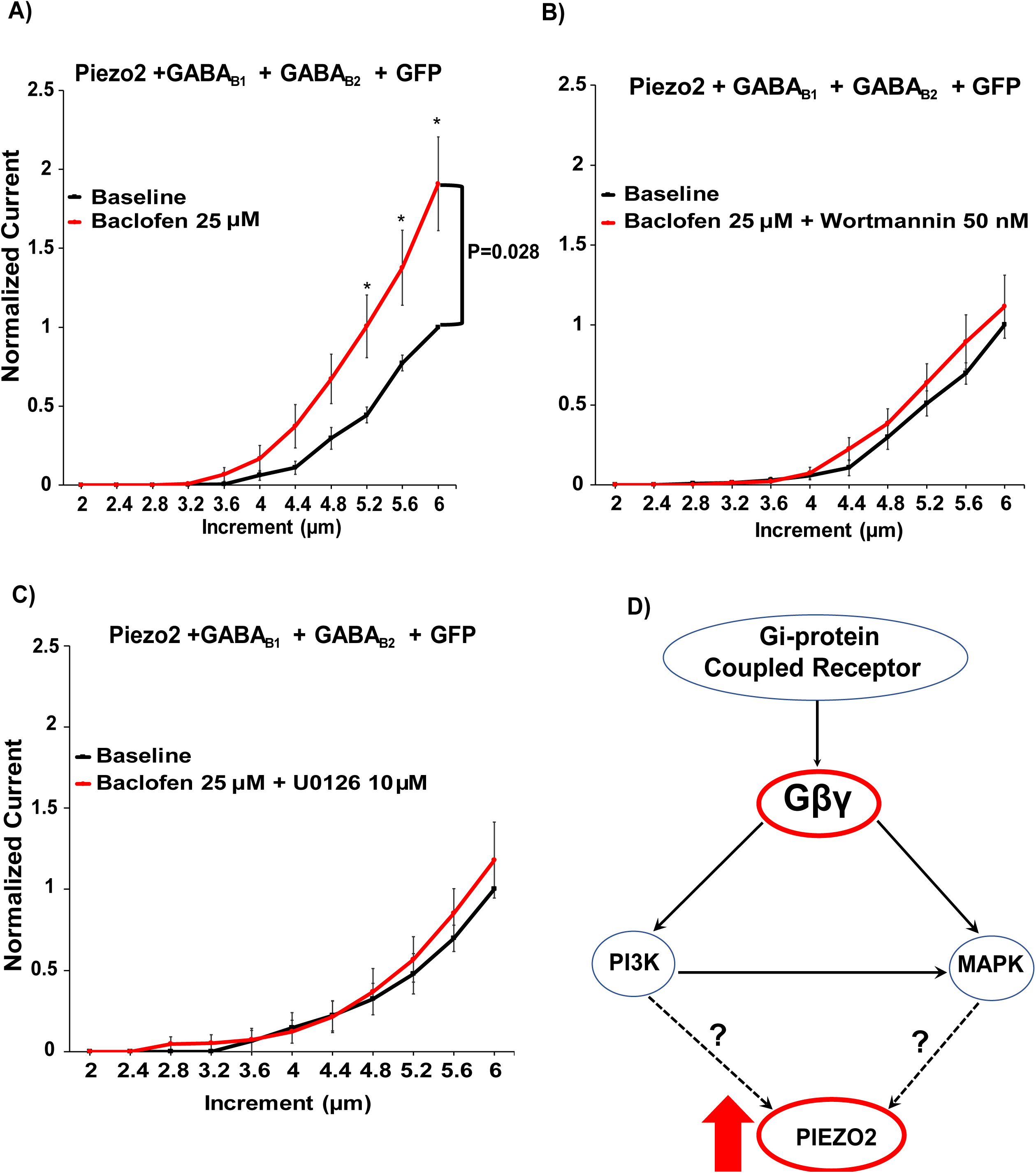
PI3K and MAPK Participate in the Potentiation of Piezo2 Currents after activation of GABA_B_ Receptors. (A-C) Quantification of normalized RA-MA Piezo2 currents in HEK293 cells transiently transfected with Piezo2^+^ GABA_B1_^+^ GABA_B2_ ^+^ GFP (A, n=12), treated with the PI3K inhibitor Wortmannin 50 nM (B, n=10) or treated with the MAPK inhibitor U0126 10 µM (C, n=7). *P<0.05, Repeated measures ANOVA with student’s t-test no corrections. Data shown as mean ±SEM. D) Prospective pathway of regulation of Piezo2 currents by Gi-coupled receptors; activation of Gi-coupled receptors promotes the release of Gβγ activating PI3K and MAPK kinase, thus indirectly potentiating Piezo2 currents.

## DISCUSSION

Here, we studied the regulation of Piezo2 channels by Gi-coupled receptors. We found that RA-MA Piezo2-like currents from large diameter DRG neurons were potentiated after activation of the Gi-coupled GABA_B_ (Fig. 1A-E) and HtrD1 receptors (Fig. 3C). However, the potentiation was not uniform amongst the DRG neurons tested because 40-50% of the neurons did not display potentiation of Piezo2-like currents when treated with baclofen (Fig. 1B and F) and sumatriptan (Fig. 3D). This phenomenon was caused probably by lack of expression of these receptors in the highly heterogenous population of DRG neurons.

To validate our findings, we co-expressed Piezo2 channels with the Gi-coupled GABA_B_ and D2 dopamine receptors in HEK293 cells and show that activation of these receptors potentiated Piezo2 currents (Fig. 2A-C; Fig. 3A). These data strongly suggest that activation of Gi-coupled receptors potentiates Piezo2 currents, and not other MA channels in DRG neurons (Lechner et al., 2009; Lechner and Lewin, 2009). Nevertheless, we cannot completely exclude the role of Gi-coupled receptors in the regulation of other MA ion channels, but the lack of identification of these proteins restrict us from studying that effect further. Our ability to reconstitute Gi-coupled receptor potentiation of Piezo2 in a heterologous system also indicates that the effect does not depend on DRG neuron specific proteins, rather it is mediated by Gi-signaling pathway components, which are present in HEK293 cells.

We also show that the activation of Gi-coupled receptors inhibits the activity of Piezo1 currents (Fig. S1A-C). This indicates that the Gi-signaling pathway is a general regulator of Piezo channels, but the regulatory mechanisms activated after receptor activation influence the activity of Piezo1 and Piezo2 differently. These findings have been in agreement with previous research showing that although Piezo1 and Piezo2 are closely related channels, they are expressed in different tissues (Coste et al., 2010), possess different kinetics of inactivation (Soattin et al., 2016), exogeneous molecules are reported to selectively activate Piezo1 channels (Syeda et al., 2015) and most likely are regulated differently by endogenous molecules.

Gi-coupled receptor activation leads to the dissociation of the trimeric G-proteins: Gαi and Gβγ. Since most of the documented effects on ion channels after activation of the Gi-signaling pathway are attributed to Gβγ (Yudin and Rohacs, 2018), we tested if the regulation of Piezo channels was mediated by Gβγ. We showed that co-expression of βARKct, which chelates Gβγ, abolished both the potentiation of Piezo2, and the inhibition of Piezo1, after activation of Gi-coupled receptors (Fig. 2F and Fig.S1E). In addition, we also showed that dialysis of Gβγ through the whole cell patch pipette also potentiated RA-MA Piezo2-like currents in DRG neurons (3E). This indicates that Gβγ was sufficient to modulate the activity of Piezo2 currents. We believe that Gαi-dependent mechanisms were unlikely to be involved in this process because Gαi is known to inhibit the activity of adenylate cyclase and activation of adenylate cyclase is required for potentiation of Piezo2 currents caused by inflammatory mediators (Dubin et al., 2012).

The potentiation of Piezo2 currents was long lasting and persisted after the removal of the Gi-coupled receptor agonist. This finding is similar to previous observations showing that activation of Gi-coupled receptors potentiated the mechanically activated two pore TREK2 K^+^channels with a very slow reversal after removal of the agonist (Lesage et al., 2000). Effects caused by direct interaction of Gβγ with ion channels are usually quickly restored after elimination of the Gi-coupled receptor agonist as shown for GIRK and TRPM3 channels (Badheka et al., 2017; Robertson and Taylor, 1986). We found no evidence for direct interaction of Gβγ with Piezo2 channels (Fig.S2), and we showed that inhibition of the Gβγ downstream effectors PI3K or MAPK abolished the potentiation of Piezo2 currents after activation of the GABA_B_ receptor (Fig. 4B and C). Both PI3K and MAPK activation has been reported to increase the activity of TRPV1 channels by increasing their surface expression (Ma et al., 2017; Stratiievska et al., 2018) but we found no evidence for increased Piezo2 trafficking to the plasma membrane in response to GABA_B_ receptor activation (Fig. S3). It was reported that PI(3,5)P_2_, a product of PI3K positively regulates Piezo2 channels (Narayanan et al., 2018). Due to lack of reliable specific probes for PI(3,5)P_2_ (Hammond et al., 2015) the involvement of this lipid in Piezo2 potentiation by Gi-coupled receptors is difficult to test, and wortmannin was reported to have only marginal effect on Piezo2 activity evoked by the knockdown of the myotubularin related protein-2 (Mtmr2), a phosphatase enzyme that dephosphorylates PI(3,5)P_2_ (Narayanan et al., 2018). Further experimentation and biochemical tools are needed to conclusively determine the mechanism of the involvement of PI3K and MAPK in regulating Piezo2 activity.

It has been well documented that the peripheral activation of Gi-coupled receptors in DRG neurons may induce hyperalgesia and promote mechanical pain (Araldi et al., 2016, 2018; Yudin and Rohacs, 2018). Intradermal hind paw injection of the migraine drug sumatriptan for example evoked both an acute hyperalgesia 30 minutes after its injection, and hyperalgesic priming, potentiating the effect of PGE2, an effect that developed several days after the injection of the drug (Araldi et al., 2016). The hyperalgesic effect of sumatriptan was proposed to play a role in migraine pain chronification in patients treated with this drug (Araldi et al., 2016). While our data showing that sumatriptan potentiated Piezo2 currents in DRG neurons may provide a potential explanation for these findings, conclusively demonstrating the involvement of Piezo2 in mechanical hypersensitivity induced by Gi-coupled receptor activation will require further studies.

In conclusion, our data showed that Gi-coupled receptors are regulators of Piezo2 and Piezo1 channels. This finding may be crucial when studying Gi-signaling pathway of MA channels in different tissues with mechanosensory properties. We also found that the potentiation and inhibition of Piezo currents could be abolished by blocking the activity of Gβγ. Additionally, our data also showed that Gβγ may not be exerting its effect directly on Piezo2 channels, but potentially by activating its downstream targets: PI3K and MAPK, suggesting a possible non-canonical mechanism of action that could serve as source of investigation for understanding mechanical hyperalgesia induced by Gi-coupled receptor activation.

## ACKNOWLEDMENTS

T.R. was supported by NIH grants NS055159 and GM093290 and J.S.D.R. was supported by NIH grant F31NS100484. The authors thank Dr. Ardem Patapoutian (Scripps Research Institute) for providing us with the Piezo1 and Piezo2 clone, Dr. Stephano Marullo (Institut Cochin) for providing the myc-GABABR1-yfp clone, Dr. Eldo Kuzhikandathil (Rutgers Brain Health Institute) for providing the D2RS-yfp clone, Dr. Diomedes Logothetis (Northeastern University) for providing the hM1 clone and Dr. Tamas Balla (NIH) for providing the Pkdc1ab-GFP (DAG sensor).

## AUTHOR CONTRIBUTIONS

J.S.D.R. and T.R. designed the experiments. J.S.D.R., Y.Y. and S.S. performed experiments and data analysis. C.H. and T.M. performed and analyzed co-immunoprecipitation experiments. J.S.D.R. and T.R. wrote and edited the manuscript. T.R. provided funding and supervised the research study.

## METHODS

### Cell Culture

#### DRG neurons

All animal procedures were approved by the Institutional Animal Care and Use Committee. Wild-type C57BL6 mice (2-to 6-month old) from either sex were anesthetized and perfused via the left ventricle with ice-cold Hank’s buffered salt solution (HBSS; Life Technologies). DRGs were harvested from all spinal segments after laminectomy and removal of the spinal column and maintained in ice-cold HBSS for the duration of the isolation. Isolated ganglia were cleaned from excess nerve tissue and incubated with type 1 collagenase (2 mg/ml; Worthington) and dispase (5 mg/ml; Sigma) in HBSS at 37°C for 30 min, followed by mechanical trituration. Digestives enzymes were then removed after centrifugation of the cells at 100g for 5 min. Isolated DRG neurons were then resuspended and seeded onto coverslips coated with a mixture of poly-D-lysine (Life Technologies) and laminin (Sigma). DRG neurons were maintained in culture in Dulbecco’s MEM/F12 (1:1) supplemented with 10% FBS (Thermo Scientific) and penicillin (100 IU/ml) and streptomycin (100 µg/ml; Life Technologies) for 16 to 48 hours before measurements.

#### HEK293 cells

HEK293 cells were obtained from the American Type Culture Collection (ATCC) (catalogue number CRL-1573, RRID:CVCL_0045) and were cultured in minimal essential medium (MEM) (Life Technologies) containing 10% (v/v) Hyclone characterized fetal bovine serum (FBS) (Thermo Scientific), and penicillin (100 IU/ml) and streptomycin (100 µg/ml; Life Technologies). Cells were transiently transfected at ∼70-80 % cell confluence with the Effectene reagent (QIAGEN) according to manufacturer’s protocol. Cells were then trypsinized and replated on poly-D-lysine-coated round coverslips 24 hours after transfection. Fluorescent cells were subjected to electrophysiological measurements after 24-48 hours. All cultured cells were kept in humidity-controlled tissue-culture incubator with 5% CO_2_ at 37°C.

### Whole-cell path clamp electrophysiology

HEK293 Cells were transiently transfected with the following cDNA constructs: mouse Piezo2 cloned into a pCMV-Sport6 vector, mouse Piezo2-GFP tagged, mouse Piezo1 cloned into a pcDNA3.1 IRES GFP vector (Coste et al., 2010), human GABA_B1_ cloned into a pCMV-Sport6 vector and GABA_B2_ cloned into a pcDNA 3.1 vector, rat βARK_CT_ cloned into a mCherry2-C1 vector, human D2RS and mouse cloned into a pcDNA 3.0 vector, Pkdc1ab-GFP (DAG sensor) cloned into a pEGFP-C1 vector and pEGFP-N1.

Whole-cell patch clamp recordings were performed at room temperature (22° to 24°C) as described previously (Yudin et al., 2011). Briefly, patch pipettes were prepared from borosilicate glass capillaries (Sutter Instrument) using a P-97 pipette puller (Sutter instrument) and had a resistance of 4-7 MΩ. After forming gigohm-resitance seals, the whole cell configuration was established and the MA currents were measured at a holding voltage of −60 mV using an Axopatch 200B amplifier (Molecular Devices). Currents were filtered at 2kHz using low-pass Bessel filter of the amplifier and digitized using a Digidata 1440 unit (Molecular Devices). All measurements were performed with extracellular solution (EC) solution containing 137 mM NaCl, 5 mM KCl, 1 mM MgCl_2_, 2 mM CaCl_2_, 10 mM Hepes and 10 mM glucose (pH adjusted to 7.4 with NaOH). The patch pipette solution contained 140 mM K^+^ gluconate, 1 mM MgCl_2_, 2 mM Na_2_ATP, 5 mM EGTA and 10 mM Hepes (pH adjusted to 7.2 with KOH). In the case of intracellular dialysis of Gβγ to cultured DRG neurons, 50 ng/ml of Gβγ or equivalent of lubrol (0.0001%) was added to the patch pipette solution.

MA currents were measured from isolated DRG neurons, or transiently transfected HEK293 cells as previously described (Borbiro et al., 2015). Briefly, Mechanical stimulation was performed using a heat-polished glass pipette (tip diameter, about 3 µm), controlled by a piezo-electric crystal drive (Physik Instrumente) positioned at 60° to the surface of the cover glass. The probe was positioned so that 10-µm movement did not visibly contact the cell but an 11.5-µm stimulus produced an observable membrane deflection. Two protocols were used to record the MA currents: *Step-increment protocol* by steadily increasing series of mechanical steps from 12 µm in 0.4-µm increments every 15 s for a stimulus duration of 200 ms and *continuous protocol* using a fixed submaximal mechanical stimulus ranging from 5.2 to 7 µm (based on the step protocols for each cell) applied every 30 s for a stimulus duration of 200 ms. Measurements from cells that showed significant swelling after repetitive mechanical stimulation were discarded (Hamill and McBride, 1997). The inactivation kinetics from MA currents were measured by fitting current decay to an exponential decay function, which measured (t) the time from the peak current and the T_inact_ (Tau-the decay constant).

### Co-Immunoprecitation

HEK293 cells on 6-well plates transfected with various constructs (indicated in figure S2) were harvested in lysis buffer (140 mM NaCl, 10 mM NAOH/HEPES, 50 mM Tris, 2.5 mM MgCl2, 1 mM EDTA, 1 mM EGTA, 0.5% Triton X-100) supplemented with protease and phosphatase inhibitor. Mouse-myc-tagged-TRPM3 cloned into bicistronic pCAGGS/IRES-GFP vector (Oberwinkler et al., 2005), mouse-myc-tagged-Piezo1 cloned into a pcDNA3.1 IRES GFP vector (Coste et al., 2015), mouse-myc-tagged-Piezo2 cloned into the Sport6 vector, and rat-myc-tagged-TRPM8 cloned into the pCDNA3 vector (Zakharian et al., 2009) were immunoprecipitated by incubating pre-cleared cell lysates with primary anti-Myc (Cell Signaling, 2276S). The immune-complex was incubated with pre-washed protein G agarose beads overnight at 4°C with gentle-rocking. Immunoprecipitants were then used for Western blotting. After six washes, precipitates were eluted from the beads by incubating at 37°C for one hour in Biorad XT loading buffer and XT reducing agent. Protein samples were run on 4–12% Bis-Tris Criterion gels and transferred to PVDF membranes. The membranes were blocked at room temperature in TBS-T with 5% milk for 1 hr and then probed overnight at 4°C with a rabbit polyclonal anti-Gβ antibody (Mirshahi et al., 2002), recognizing Gβ1, Gβ2, Gβ3 Gβ4 (T-20, SC-378, Santa Cruz) diluted 1:1000 in TBS-T with 5% milk. Secondary antibody used was donkey-anti-rabbit HRP (Thermo-Fisher, A16035) 1:5000 in 5% Milk. All blots were processed with SuperSignal West Pico Chemiluminescent Substrate (Thermo Fisher Scientific, Waltham, MA) and imaged with a Fuji Imager.

### Total Internal Reflection Fluorescence (TIRF) Imaging

TIRF measurements were performed as previously described (Liu et al., 2019). Briefly, an Olympus IX-81 inverted microscope was equipped with an ORCA-FLASH 4.0 camera (Hamamatsu) and a single line TIRF illumination module. Excitation light was provided by 488 nm argon laser through a 60x NA 1.49 TIRF objective. For experiments in Fig.S3, HEK293 cells were transfected with Piezo2-GFP tagged, Piezo2-GFP tagged with GABA_B_Rs or human muscarinic 1 (M1) receptors with Pkdc1ab-GFP tagged (DAG sensor). Cells were plated on 25 mm glass coverslips, and solution were manually applied to the chamber with gentle mixing. Data was analyzed using Metamorph and Image J software.

### Statistics

Data analysis was performed in Excel and SPSS. Data collection was randomized. Data in all measurements were normalized by dividing every current value by the highest current value recorded from the baseline measurements of each cell before baclofen application. The normality of the data was verified with Shapiro-Wilk test. Data were analyzed with t-test, or Repeated-measures Analysis of Variance (ANOVA) with Student’s t-test no corrections (Jia et al., 2013; Saville, 2003), p values are reported in the figure legends. Data are plotted as mean ^+^/-standard error of the mean (SEM) for most experiments.

---

**Figure S1.**
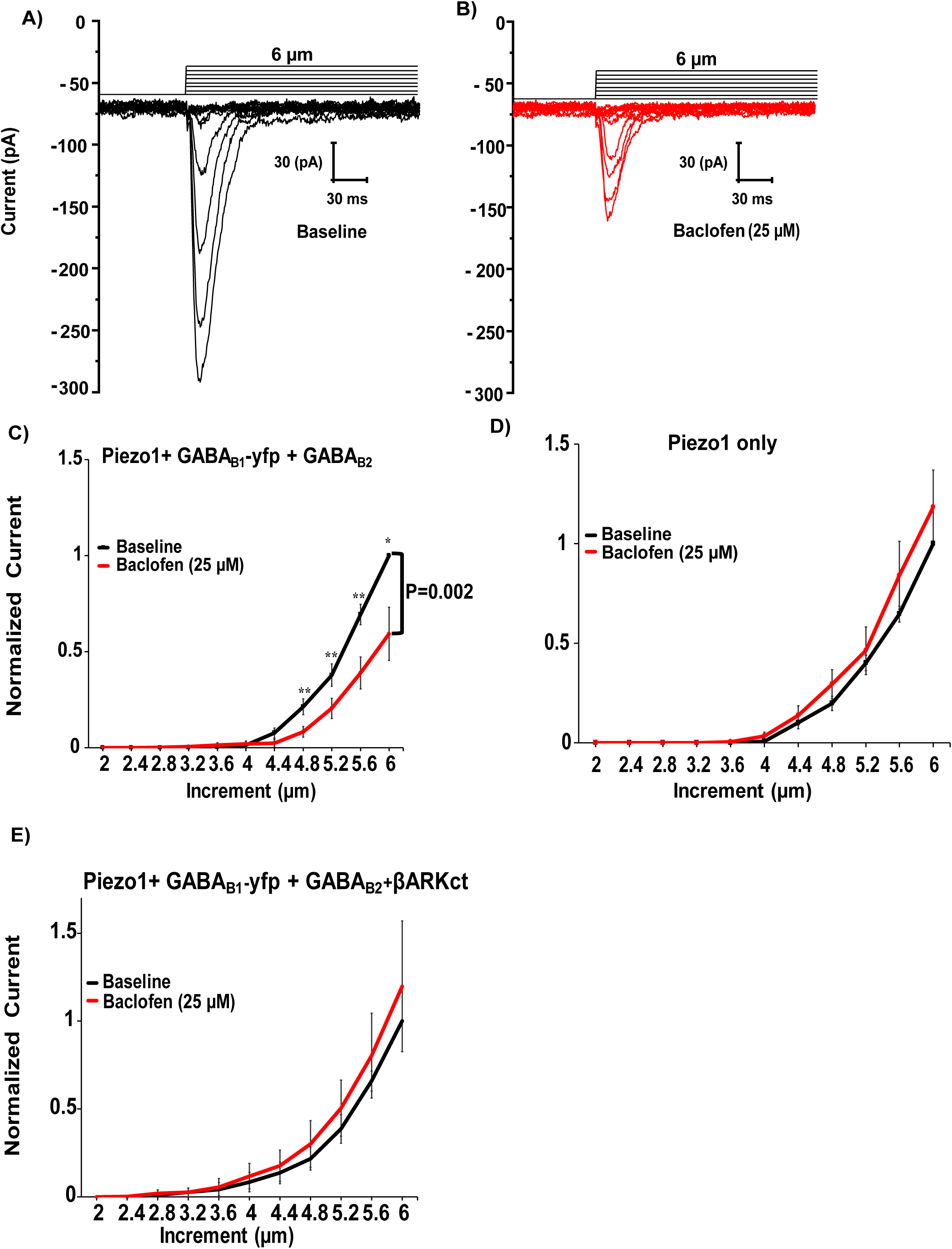
Activation of GABA_B_ Receptors Inhibits Piezo1 Currents in Heterologous Systems via Gβγ. Related to figure 2. (A-B) Representative of RA-MA currents recorded in whole cell patch clamp experiments at −60 mV in HEK293 cells transiently transfected with Piezo1^+^GABA_B1_-yfp^+^ GABA_B2_ in response to mechanical stimulation with a blunt glass probe displaced 2-5.6 µm every 15 s by the repetitive mechanical stimuli before (black) and after (red) exposure to 25 µM baclofen. The mechanical step protocol that indicates the displacement of mechanical probe is also shown. (C-E) Quantification of normalized MA currents of HEK293 cells transfected with Piezo1^+^ myc-GABA_B1_-yfp ^+^ GABA_B2_ (C, n=13), with Piezo1-IRES-eGFP (D, n=9) and Piezo1^+^ GABA_B1_-yfp ^+^ GABA_B2_ ^+^ βARKct (E, n=8) before and after application of 25 µM baclofen (*P<0.05, **P<0.01, Repeated measures ANOVA with student’s t-test no corrections). Data shown as mean ±SEM.

**Figure S2:**
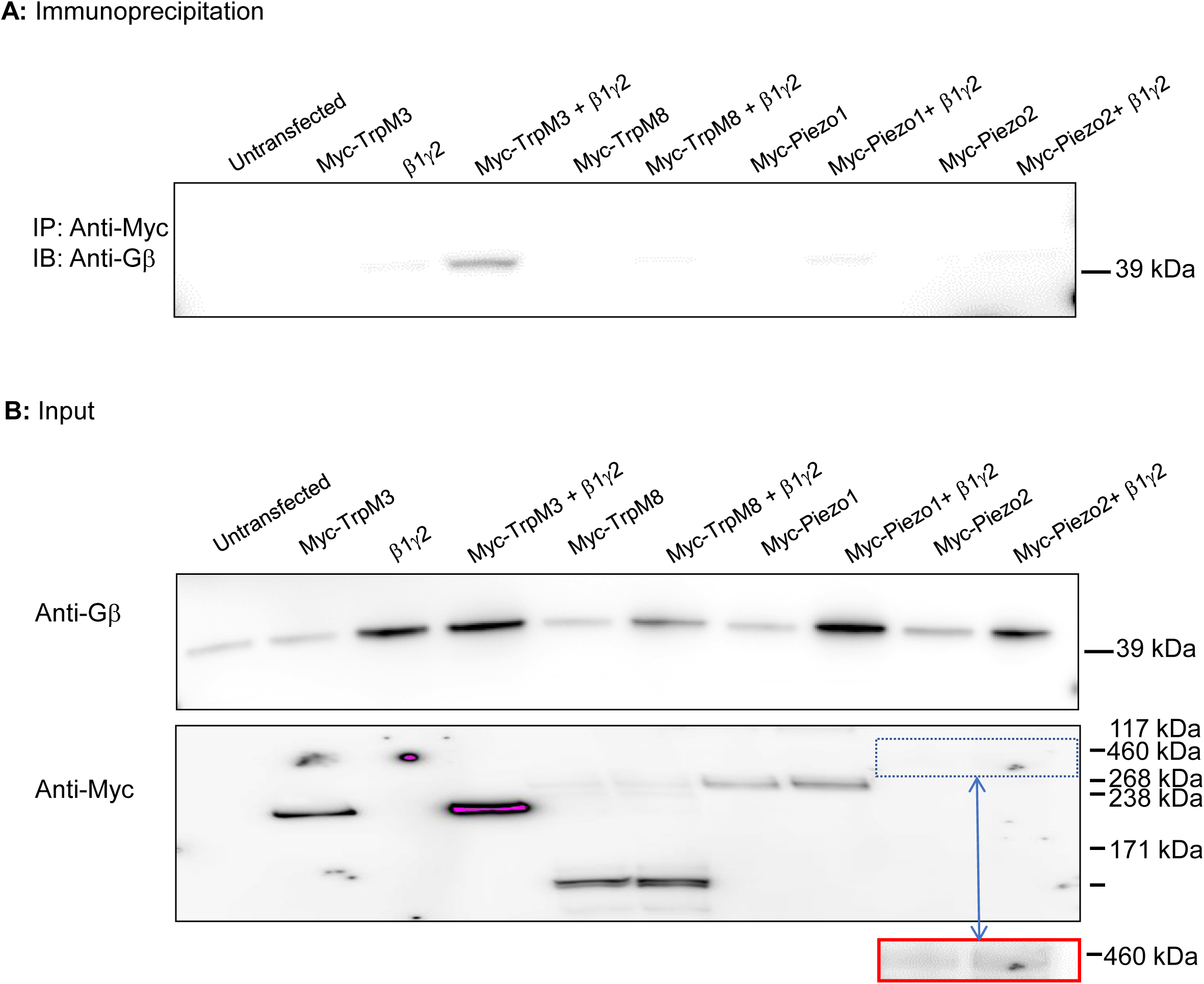
Gβγ do not interact with Piezo1 and Piezo2. Related to figure 4. (A) Myc-tagged Piezo1 and Piezo2, as well as Myc-TRPM3 (positive control) and Myc-TRPM8 (negative control) were co-expressed with or without exogenous Gβ1γ2 in HEK293 cells. Lysates were immunoprecipitated with anti-Myc and immunoblotted with anti-Gβ. Gβ could only be immunoprecipitated with Myc-TrpM3, but none of the other My-tagged channels. *A very weak band is seen in all lanes expressing exogenous Gβγ which we believe is due to a small non-specific anti-body interaction*. Data are representative of 3 replicates. (B) Inputs from the lysate used for immunoprecipitation show expression of all the transfected constructs as well as native Gβ. Piezo2 routinely showed low expression, but could be detected at higher exposures (red outlined inset)

**Figure S3.**
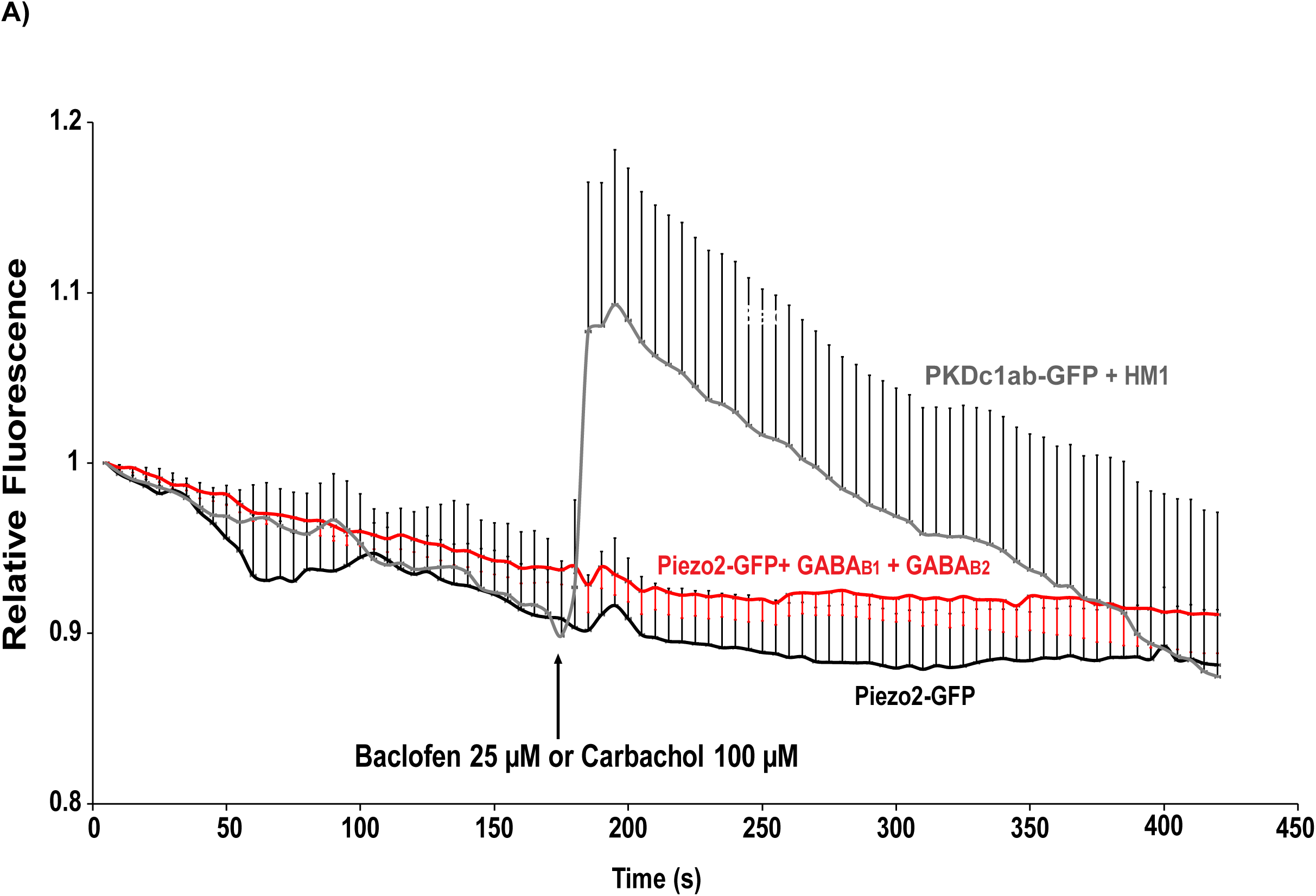
Activation of GABAB Receptors does not Increase the Proportion of Piezo2 Channels at the Plasma Membrane. Related to figure 4. (A) Normalized time course of changes in fluorescence intensity from HEK293 cells transiently transfected with either Piezo2-GFP (black), Piezo2-GFP^+^ GABA_B1_^+^ GABA_B2_ (red) or muscarinic acetylcholine receptor 1 (HM1) ^+^ PKDc1ab-GFP (gray line-a DAG sensor used as positive control) using TIRF microscopy. Transfected HEK293 cells were treated for 7 mins with 25 µM baclofen or 100 µM carbachol (positive control). There was not statistical change in fluorescence intensity in baclofen-treated groups. Data were normalized to the mean intensity values at the beginning of the recording prior to baclofen or carbachol application. Data shown as mean ±SEM

